# Protective mucosal immunity against SARS-CoV-2 after heterologous systemic RNA-mucosal adenoviral vector immunization

**DOI:** 10.1101/2021.08.03.454858

**Authors:** Dennis Lapuente, Jana Fuchs, Jonas Willar, Ana V Antão, Valentina Eberlein, Nadja Uhlig, Leila Issmail, Anna Schmidt, Friederike Oltmanns, Antonia Sophia Peter, Sandra Mueller-Schmucker, Pascal Irrgang, Kirsten Fraedrich, Andrea Cara, Markus Hoffmann, Stefan Pöhlmann, Armin Ensser, Cordula Pertl, Torsten Willert, Christian Thirion, Thomas Grunwald, Klaus Überla, Matthias Tenbusch

## Abstract

Several effective SARS-CoV-2 vaccines are currently in use, but in the light of waning immunity and the emergence of novel variants, effective boost modalities are needed in order to maintain or even increase immunity. Here we report that intranasal vaccinations with adenovirus 5 and 19a vectored vaccines following a systemic DNA or mRNA priming result in strong systemic and mucosal immunity in mice. In contrast to two intramuscular injections with an mRNA vaccine, the mucosal boost with adenoviral vectors induced high levels of IgA and tissue-resident memory T cells in the respiratory tract. Mucosal neutralization of virus variants of concern was also enhanced by the intranasal boosts. Importantly, priming with mRNA provoked a more comprehensive T cell response consisting of circulating and tissue-resident memory T cells after the boost, while a DNA priming induced mostly mucosal T cells. Concomitantly, the intranasal boost strategies provided protection against symptomatic disease. Therefore, a mucosal booster immunization after mRNA priming is a promising approach to establish mucosal immunity in addition to systemic responses.

## Introduction

The severe acute respiratory syndrome coronavirus 2 (SARS-CoV-2) emerged in late 2019 and caused a worldwide pandemic accounting for over 190 million infections and 4 million deaths at the time of this report ^1^. In an unprecedented speed, academic institutions and biotech companies developed, evaluated, and licensed several SARS-CoV-2 vaccines. Beside traditional approaches like protein subunit or inactivated virus vaccines, gene-based vaccines were at the forefront of the developmental process and the first to become licensed ^2^.

Vaccines based on messenger RNA (mRNA) or adenoviral vectors (Ad) demonstrated efficacy against SARS-CoV-2 infections and, most importantly, against severe coronavirus disease 2019 (COVID-19) and death ^3–6^. Humoral as well as cellular immune responses against the spike (S) surface protein were successfully induced by both types of vaccines ^7–11^. However, breakthrough infections of fully vaccinated individuals have been reported and the numbers might increase in the phase of waning immunity ^12–18^. The impact of immune escape and newly emerging virus variants (i.e. variants of concern, VOCs) is controversially discussed in some of these studies. Upon breakthrough infections, virus replication in the respiratory tract is approximately four- to six-fold reduced compared to unvaccinated and virus shedding seems to be shorter in duration ^14, 19^. Importantly, Public Health England reported that after the first dose of an mRNA (Comirnaty®) or viral vector vaccine (Vaxzevria®) the likelihood of household transmission drops by 40-50% ^20^. On one hand, these observations underline that the current vaccination campaigns can end the pandemic phase by reducing the basic reproduction number below 1. On the other hand, however, it also demonstrates that transmission is still possible by vaccinated individuals posing a risk to vulnerable communities.

While the currently approved vaccines induce systemic immune responses, they probably do not evoke mucosal immunity in form of mucosal, secretory immunoglobulin A (IgA) or tissue-resident memory T cells (T_RM_). Secretory, polymeric IgA can neutralize incoming viral particles at the mucosal surface before infection of epithelial cells takes place, which is important for an optimal protection against respiratory virus infections ^21–23^. Furthermore, IgA enables specific effector functions by cross-linking the Fcα-receptor, and polymeric forms of IgA might increase antibody avidity ^24^. So far, the only licenced intranasal vaccines are live-attenuated influenza vaccines (LAIV). Nasal IgA contributes to the efficacy of LAIV in children ^25^ and also correlates with protection in experimental human challenge studies ^26^. Importantly, local antigen deposition by mucosal vaccination routes is key for an induction of mucosal IgA as shown in humans ^24, 27–29^ and animal models ^30–32^. While IgA can be effectively induced by intranasal delivery of protein-based vaccines, an efficient induction of respiratory CD8^+^ T_RM_ usually requires local antigen production in the mucosa followed by major histocompatibility complex-I-mediated peptide-presentation by stromal and, most importantly, by migratory CD103^+^ dendritic cells^33^. CD8^+^ T_RM_ localize within the respiratory epithelium or the airways and can respond immediately in case of secondary infections. In contrast to circulatory T cell memory phenotypes like central memory (T_CM_), effector-memory (T_EM_), or effector T cells (T_EFF_), T_RM_ do not significantly recirculate ^34, 35^. Thus, one feature of T_RM_ is the direct localization at barrier tissues, which makes a time-consuming migration into the inflamed lung redundant. A second remarkable characteristic is the ability to exert innate and adaptive functions within a few hours after secondary infection ^36, 37^, in part due to the storage of ready-made mRNAs encoding cytokines like IFNγ at steady state ^38, 39^. Altogether, these unique features of mucosal immune responses enable an immediate and effective countermeasure against pulmonary infections as described for flu ^40, 41^, respiratory syncytial virus (RSV) ^42^, and *Mycobacterium tuberculosis* ^43, 44^. The great majority of these finding were generated in animal models, partly due to the invasive nature of bronchoalveolar lavages (BAL) and biopsy sampling. However, small experimental human challenge studies started to look precisely at the role of mucosal immunity against respiratory viruses^45, 46^.

A few preclinical studies investigated intranasal SARS-CoV-2 vaccines so far. In a series of publications, one group reported protective efficacy of a one shot vaccination with an chimpanzee adenoviral vector (ChAd) vaccine encoding for the spike protein in mice, hamsters, and rhesus macaques ^47–49^. Importantly, van Doremalen et al. have shown that intramuscular ChAd vaccination prevents pneumonia in macaques but allow for virus replication in the upper respiratory tract ^50^. However, administered intranasally, the vaccine attenuated nasal virus replication more efficiently ^51^. It is important to investigate intranasal vaccine candidates not only as standalone modality but also in the context of pre-existing immunity induced by a previous vaccination. On one hand, this is important due to the broad employment SARS-CoV-2 vaccines in recent vaccination campaigns. On the other hand, first clinical data point towards suboptimal immunogenicity of solely intranasal vaccinations against SARS-CoV-2 in humans without pre-existing immunity, but also provides evidence for robust immunity after heterologous prime-boost vaccinations ^52, 53^.

Here we demonstrate that a systemic DNA or mRNA prime followed by an intranasal boost with an adenoviral serotype 5 vector (Ad5) enables a comprehensive systemic and local T cell immunity as well as substantial mucosal neutralization of SARS-CoV-2 VOCs. Concomitantly, the mucosal boost strategies led to an efficient control of virus replication after experimental infection comparable to homologous systemic immunizations.

## Results

### A systemic DNA prime significantly increases the mucosal immunogenicity of an intranasal adenoviral vector vaccine

In this first part of our study, we evaluated the immunogenicity of mucosally applied viral vector vaccines as a single shot vaccine or as a booster after an intramuscular DNA prime immunization. To this end, codon-optimized sequences encoding the full-length S and nucleocapsid (N) proteins of SARS-CoV-2 were inserted into pVax-1 expression plasmids and into replication-deficient adenoviral vector vaccines based on serotype 5 (Ad5) or serotype 19a (Ad19a). BALB/c mice were immunized intranasally with the Ad5- or Ad-19a-based vaccines either without prior treatment or four weeks after an intramuscular DNA immunization with S- and N-encoding plasmids (Fig. 1 A). Two weeks later, SARS-CoV-2 specific antibody responses were analysed in serum and BAL, whereas the local and systemic T cell responses were determined in lungs and spleens, respectively.

**Figure 1:**
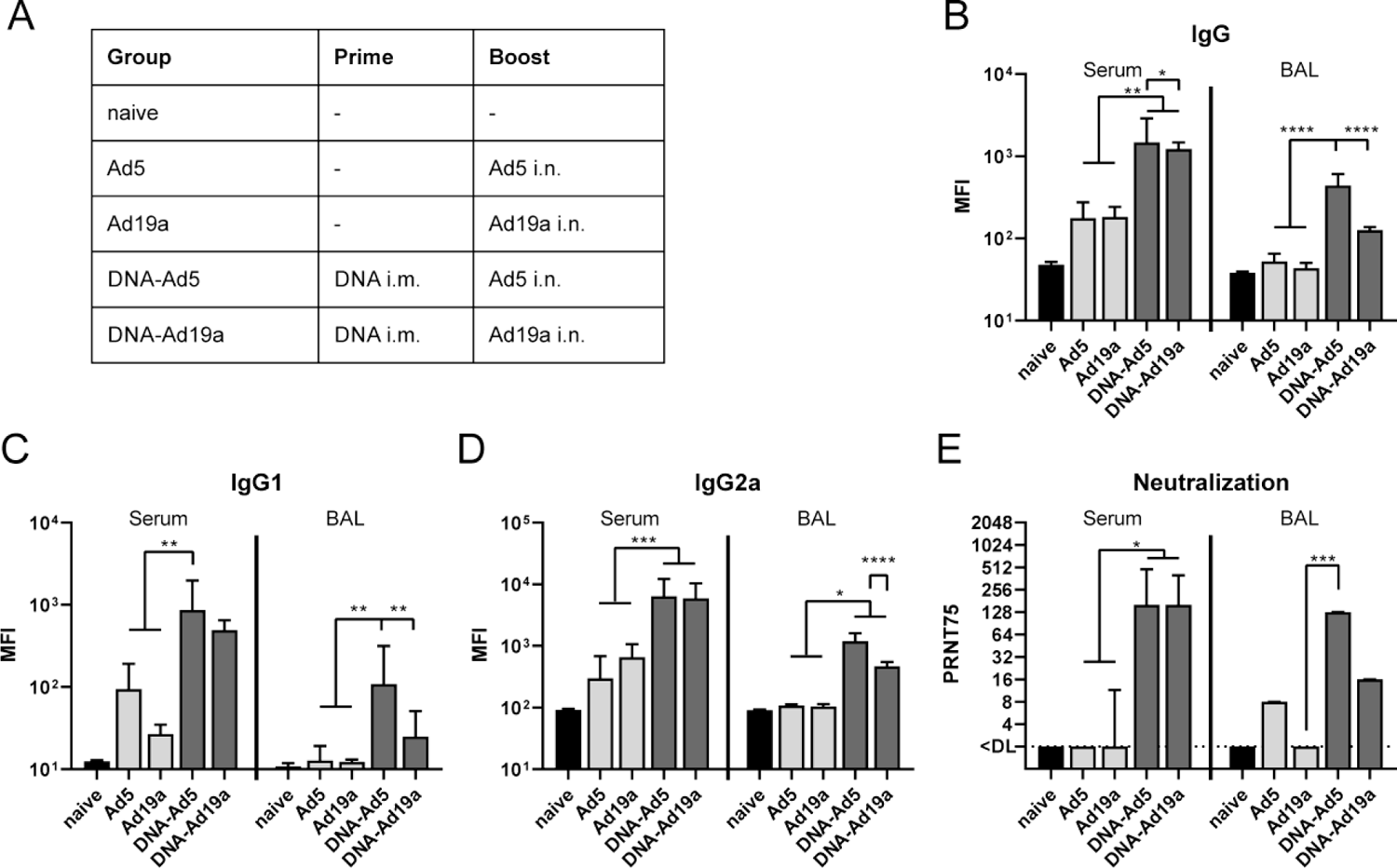
Humoral responses after intranasal immunization with Ad5- or Ad19a-based viral vector vaccines. (A) BALB/c mice were immunized intranasally with Ad5- or Ad19a-based vectors encoding the N and S protein of SARS-CoV-2 (2×10^6^ infectious units per vector). Mice from the heterologous prime-boost groups were primed four weeks before by intramuscular injection of N- and S-encoding DNA plasmids (10 µg per plasmid) followed by electroporation. Serum antibody responses were analysed thirteen days and mucosal immune responses in the BALs fourteen days after the mucosal immunization. Spike-specific IgG (B), IgG1 (C), and IgG2a (D) were assessed by a flow cytometric approach (dilutions: sera 1:400, BAL 1:100). Plaque reduction neutralization titres (PRNT75) were determined by in vitro neutralization assays (B). Bars represent group medians with interquartile ranges; naïve n=4; DNA-Ad5 n=5; other groups n=6. Data were analysed by one-way ANOVA followed by Tukey’s post test (B-D) or by Kruskal-Wallis test (one-way ANOVA) followed by Dunn’s multiple comparison test (E). Statistically significant differences were indicated only among the different vaccine groups (*, p<0.05; **, p<0.005; ***, p<0.0005; ****, p<0.0001).

In our flow cytometric assay ^54^, spike-specific IgG, IgG1, and IgG2a could be easily detected in serum and BAL of animals treated with the prime-boost strategies, while antibodies in the BAL after a single dose of Ad19a or Ad5 were almost absent (Fig. 1 B-D). Comparing the two adenoviral vectors as booster vaccines, the serotype 5 induced significant higher levels of S-specific antibodies in the BAL, although the antibody levels in the sera were comparable for the both groups. A predominant polarisation of the antibody response towards either IgG1 or IgG2 is not indicated for any vaccine group (Fig. 1 C and D). Similar trends were also observed for N-specific antibody levels in sera and BALs (Fig. S1). In line with the amount of S-binding antibodies, profound virus neutralization was detected in sera and BAL samples from the groups DNA-Ad5 and DNA-Ad19a, while Ad5 or Ad19a alone did not induce significant levels of neutralizing antibodies (Fig. 1 E). Given the differences in the local antibody levels, the IgA response in the BAL towards specific domains of the S protein were analysed in more detail by ELISA (Fig. 2 A-C). These results confirmed that intranasal applications of Ad5-based vectors induce higher S-specific IgA levels than Ad19a-based vectors and these responses benefit from a systemic DNA prime. Furthermore, the vaccine-induced antibodies were directed against S1 including the receptor-binding domain (RBD) as well as against the S2 domain of the spike protein (Fig. 2 A-C).

**Figure 2:**
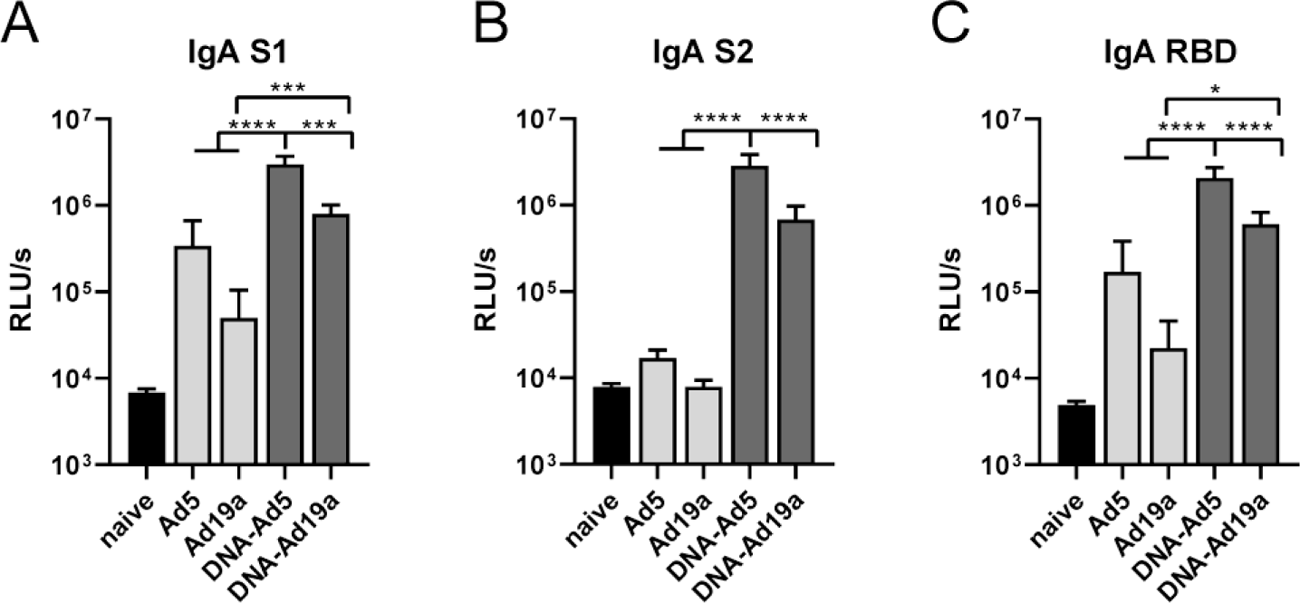
Mucosal, Spike-specific IgA responses. BALB/c mice were vaccinated according to Fig. 1 A. BAL samples were tested for spike-specific IgA directed against the domains of S1 (A), S2 (B), or RBD (C) by ELISA (dilution: 1:10). Bars represent group medians with interquartile ranges; naïve n=4; DNA-Ad5 n=5; other groups n=6. Data were analysed by one-way ANOVA followed by Tukey’s post test. Statistically significant differences were indicated only among the different vaccine groups (*, p<0.05; **, p<0.005; ***, p<0.0005; ****, p<0.0001).

Next, we assessed the induction of cellular immune responses in the lung by the different vaccination schemes. Intravascular staining (iv-labeling) ^55^ was used to differentiate between circulating T cells present in the lung endothelium during sampling (iv-) and T_RM_ (iv+). Since specific MHC-I multimers were not available at the time of this study, antigen-experienced cells were identified by the expression of CD44 (gating strategy shown in Fig. S2). Similar to the humoral responses, CD44^+^ CD8^+^ T cells in the lung were most efficiently induced by the DNA-Ad5 scheme, although all vaccinated animals mounted vaccine-induced cellular responses (Fig. 3 A). The vast majority of lung CD8^+^ T cells were protected from the iv-labeling in all groups, and the most prominent T_RM_ phenotype was CD103^+^CD69^+^ (Fig. 3 B). Antigen-specific CD4^+^ and CD8^+^ T cells were identified by ex vivo restimulation with peptide pools covering major parts of S and the complete N protein, respectively, followed by intracellular staining of accumulated cytokines (gating strategy in Fig. S3). The highest percentages of S-reactive CD8^+^ T cells were detected in the lungs of DNA-Ad5 treated animals with the majority of them predominantly producing IFNγ (Fig. 4 A). Substantial numbers of this cell population were also present after the single shot vaccination with Ad5 reaching comparable levels to the ones of DNA-Ad19a treated animals. Differences in the percentages of CD8^+^ T cells expressing IL-2 or TNFα were less pronounced, and polyfunctional T cells positive for all four analytes including the degranulation marker CD107a were rarely found in all animals. In contrast, significantly elevated percentages of CD8^+^ T cells producing IFNγ or TNFα as well as polyfunctional CD8^+^ T cells were detected in the spleens of DNA-Ad19a treated animals (Fig. 4 C). Albeit at overall lower frequencies, the same observation was made for N-reactive CD8^+^ T cells in lungs and spleens (Fig. S4 A and C). Strong S- and N-specific CD4^+^ T cell responses were detected in all animals that received a prime-boost vaccination (Fig. 4 and Fig. S4). In contrast to the CD8^+^ T cells, the majority of the CD4^+^ T cells were polyfunctional indicated by the simultaneous expression of IFNγ, TNFα and IL-2. Again, immunization with the Ad19a-based vectors resulted in higher systemic responses measured in the spleen, whereas the mucosal response in the lung was more pronounced after delivery of Ad5-based vectors (Fig. 4 C and D, Fig. S4 C and D).

**Figure 3:**
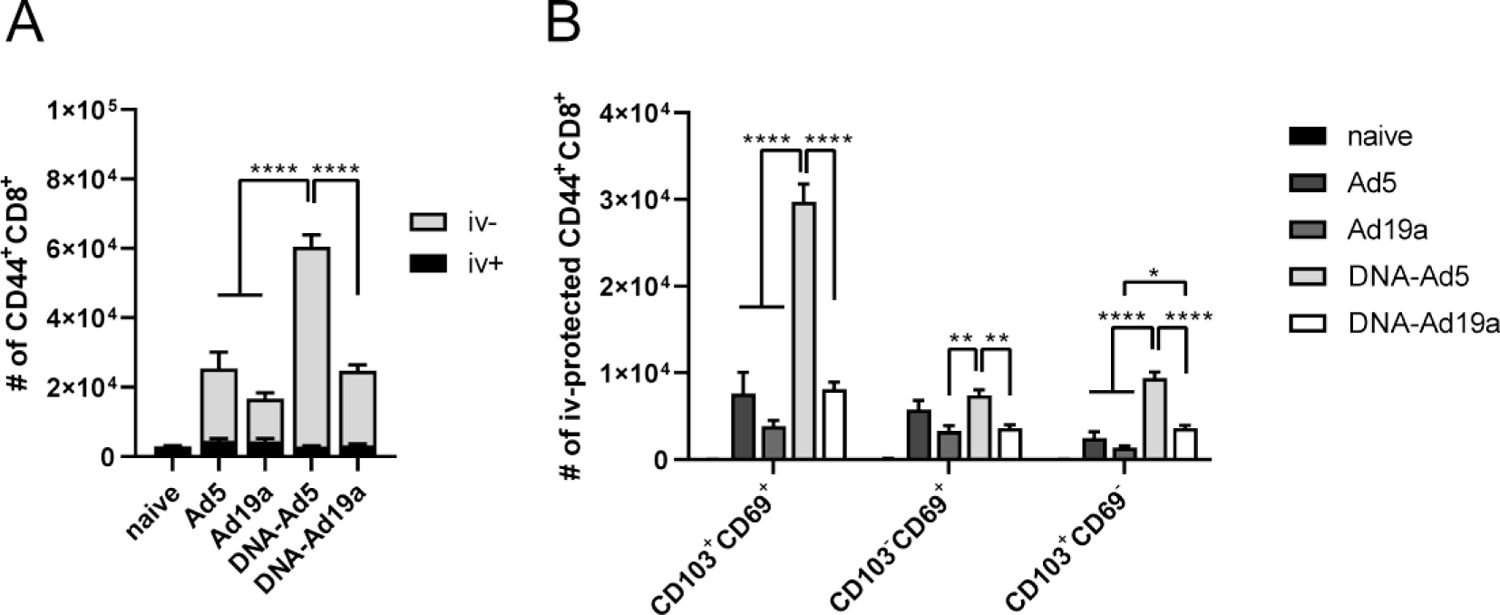
Tissue-resident memory T cell subsets in the lung. BALB/c mice were vaccinated according to Fig. 1 A. In absence of suitable MHC-I multimers, antigen-experienced CD8^+^ T cells were identified by CD44 staining (A). Intravascular staining was used to differentiate between circulating (iv+) and tissue-resident (iv-) memory cells. Tissue-resident phenotypes were assessed by staining for CD69 and/or CD103 within the iv-protected memory compartment (B). The gating strategy is shown in figure S2. Bars represent group means with SEM; naïve n=4; DNA-Ad5 n=5; other groups n=6. Data were analysed by one-way ANOVA followed by Tukey’s multiple comparison test. Statistical significant differences were indicated only among the different the vaccine groups (*, p<0.05; **, p<0.005; ***, p<0.0005; ****, p<0.0001) and in case of (A) for the total CD44 population including iv-protected and iv-labelled.

**Figure 4:**
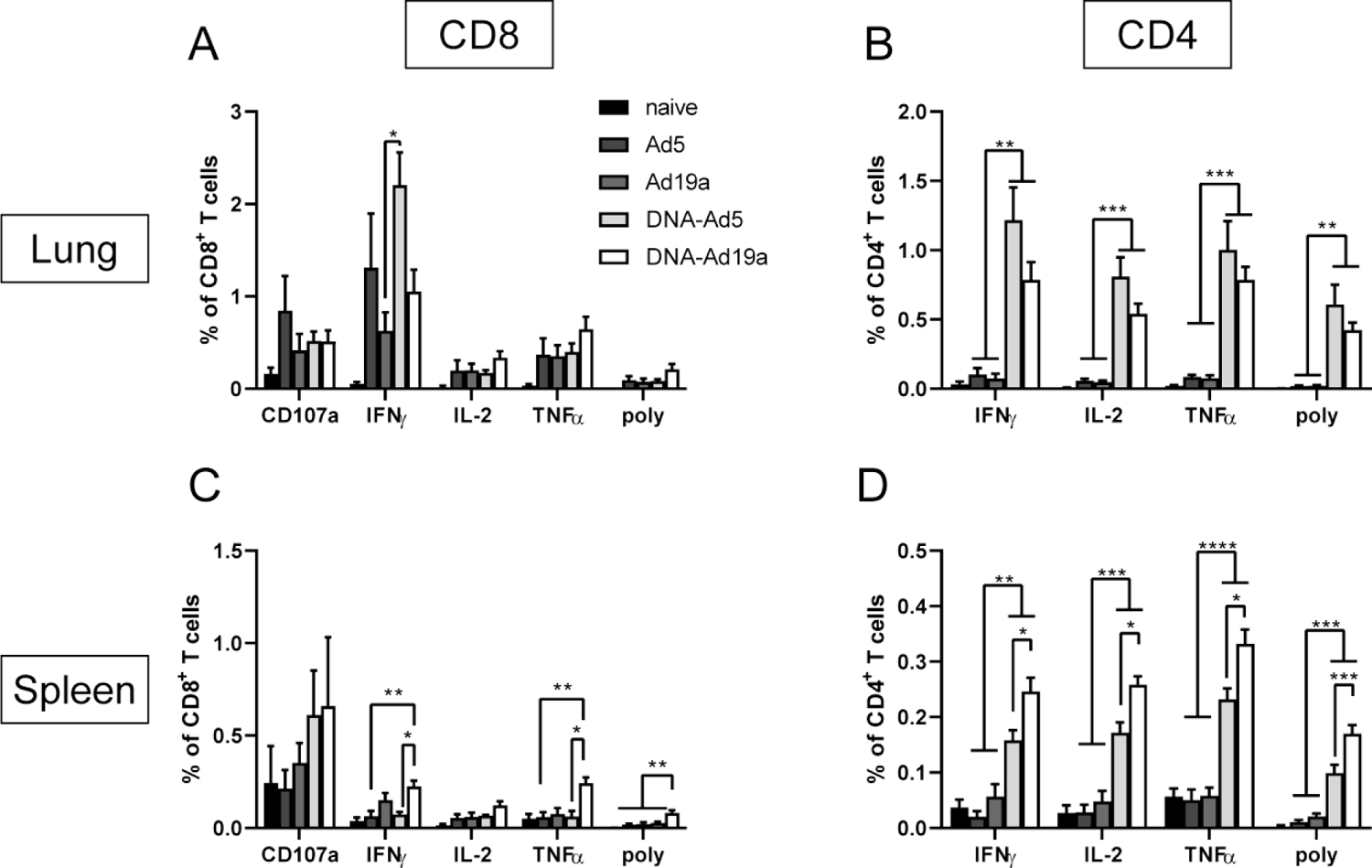
Spike-specific T cell responses after intranasal immunization with Ad5- or Ad19a-based viral vector vaccines. BALB/c mice were vaccinated according to Fig. 1 A. Lung and spleen homogenates were restimulated with peptide pools covering major parts of S and the responding CD8^+^ (A and C) and CD4^+^ T cells (B and D) were identified by intracellular staining for accumulated cytokines or staining for CD107a as degranulation marker. The gating strategy is shown in figure S3. Bars represent group means with SEM; naïve n=4; DNA-Ad5 n=5; other groups n=6. Data were analysed by one-way ANOVA followed by Tukey’s multiple comparison test. Statistically significant differences were indicated only among the different vaccine groups (*, p<0.05; **, p<0.005; ***, p<0.0005; ****, p<0.0001). poly; polyfunctional T cell population positive for all assessed markers.

Taken together, Ad5 proved a higher immunogenicity as mucosal vaccine vector compared to Ad19a and resulted in strong cellular and humoral immune responses against SARS-CoV-2 antigens if combined with an intramuscular DNA prime immunization.

### An intranasal boost following mRNA vaccination potentiates mucosal antibody responses with pronounced neutralization breadth

Since mRNA vaccines are currently in use for mass vaccination campaigns in many countries, we wanted to compare the differential effects of a DNA or mRNA prime on the immunogenicity of a mucosal booster. Therefore, the previously described DNA-Ad5 scheme was compared to an mRNA prime (Comirnaty®, Biontech/Pfizer) followed by an intranasal Ad5 boost (RNA-Ad5). Moreover, two vaccine groups that received two intramuscular injections with either mRNA (2x RNA) or an adenoviral vector (2x Ad5) reflecting current SARS-CoV-2 vaccination strategies were included (Fig. 5 A). These experiments were performed in C57BL/6 mice to allow correlations to efficacy data in K18-hACE2 mice.

**Figure 5:**
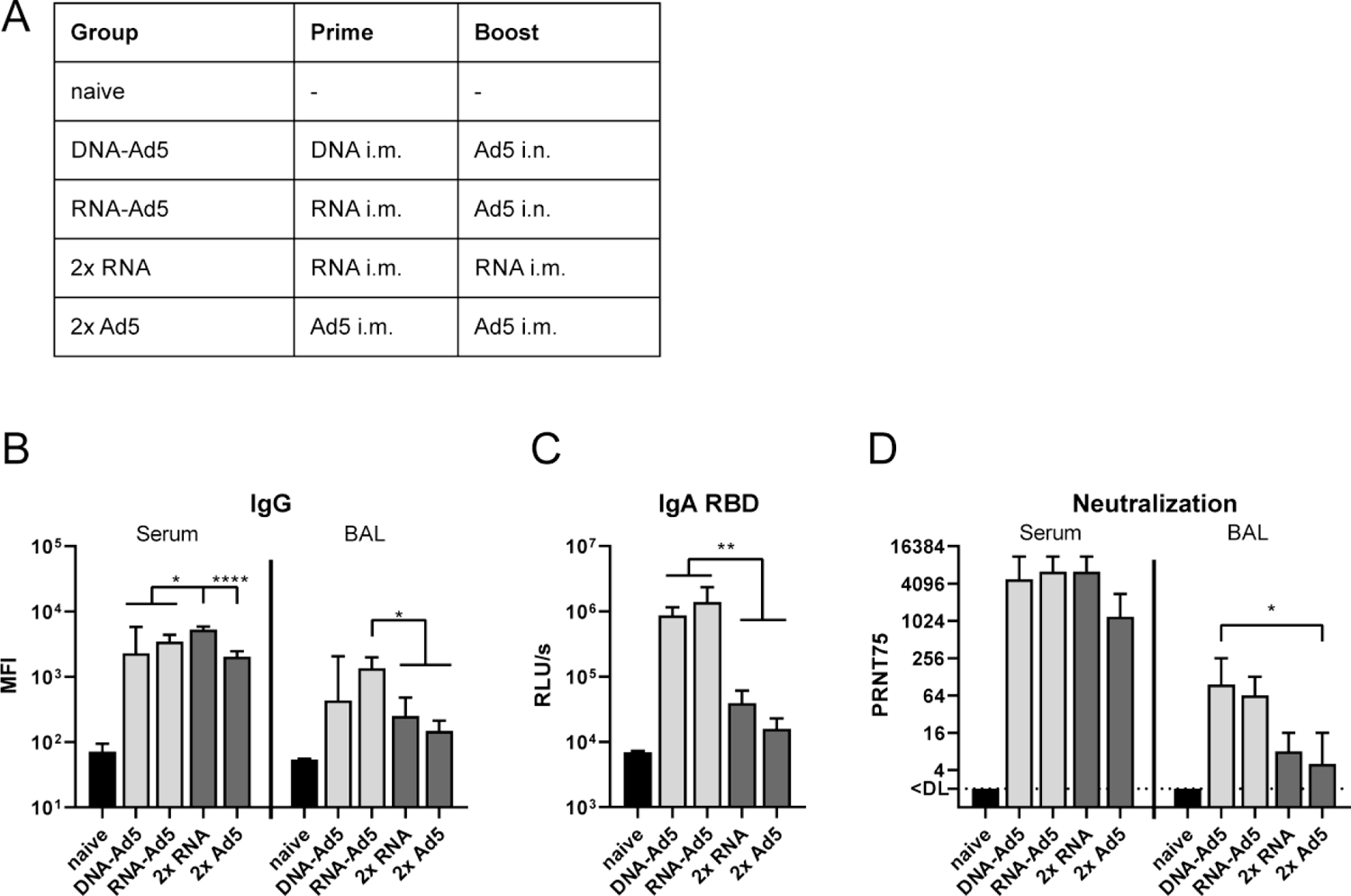
Humoral responses after homologous or heterologous prime-boost vaccination. (A) C57BL/6 mice received an intramuscular prime immunization with the spike-encoding DNA (10 µg), Ad5-S (10^7^ infectious units), or the mRNA vaccine, Comirnaty® (1 µg). Mice from the heterologous prime-boost groups were boosted four weeks later intranasally with Ad5-S (10^7^ infectious units). The homologous prime-boost groups received a second dose of mRNA (1 µg) or Ad5-S (10^7^ infectious units) intramuscularly. Serum antibody responses were analysed 21 days and mucosal immune responses four weeks after the boost immunizations. Spike-specific IgG (B) were assessed by a flow cytometric approach (dilutions: Sera 1:800, BAL 1:20). BAL samples were tested for spike-specific IgA directed against RBD by ELISA (C). Plaque reduction neutralization titres (PRNT75) were determined by in vitro neutralization assays (D). Bars represent group medians with interquartile ranges; sera all groups n=8; BALs RNA-Ad5 n=7, other groups n=8. Data were analysed by one-way ANOVA followed by Tukey’s post test (B and C) or Kruskal-Wallis test (one-way ANOVA) followed by Dunn’s multiple comparison test (D). Statistically significant differences were indicated only among the different vaccine groups (*, p<0.05; **, p<0.005; ***, p<0.0005).

Four weeks after the boost immunization, all vaccinated animals reached high levels of anti-S IgG in the serum (Fig. 5 B and Fig. S5). However, the anti-S IgG levels after the homologous RNA vaccination were significantly higher than in all other groups. Interestingly, this order does not reflect the anti-S response measured four weeks after the prime immunization. Here, the intramuscular injection of Ad5 induced the highest antibody levels, most probably by inducing more potent IgG2a responses than the RNA vaccine (Fig. S6). Contrary, the IgG levels detected in BALs were higher in the groups receiving the intranasal Ad5 boost vaccination (Fig. 5 B). Additionally, significantly increased local IgA antibody levels could be detected for both groups in a RBD-specific ELISA (Fig. 5 C). On a functional level, the higher amounts of RBD-specific antibodies were mirrored by higher neutralizing capacities in the BALs of the groups DNA-Ad5 or RNA-Ad5 (Fig. 5 D). Interestingly, the high amount of neutralizing antibodies in the sera were not significantly different among the vaccine groups independent of the route of the boost immunization.

Since mucosal antibodies might be most important for preventing an initial infection and thereby transmission, we evaluated the protective capacity against SARS-CoV-2 VOCs in pseudotype-based virus neutralization assays (Fig. 6). Here, the most robust and broadest responses were detected in the BALs of RNA-Ad5 treated animals with decreasing neutralizing potencies against spike proteins from SARS-CoV-2 lineages B.1.1.7 (alpha variant)/P.1 (gamma variant) to B.1.351 (beta variant), and finally B.1.617.2 (delta variant). Interestingly, the DNA-Ad5 scheme resulted in comparable IC75 titres against alpha and delta as RNA-Ad5, but was less potent against the beta variant. This might reflect the different nature of the encoded S protein sequences. Finally, the solely systemic vaccination schedules provoked 4 - 32-fold lower titres of mucosal neutralization against alpha, beta, and gamma, whereas no neutralization of delta spike-pseudotyped reporter virus could be observed.

**Figure 6:**
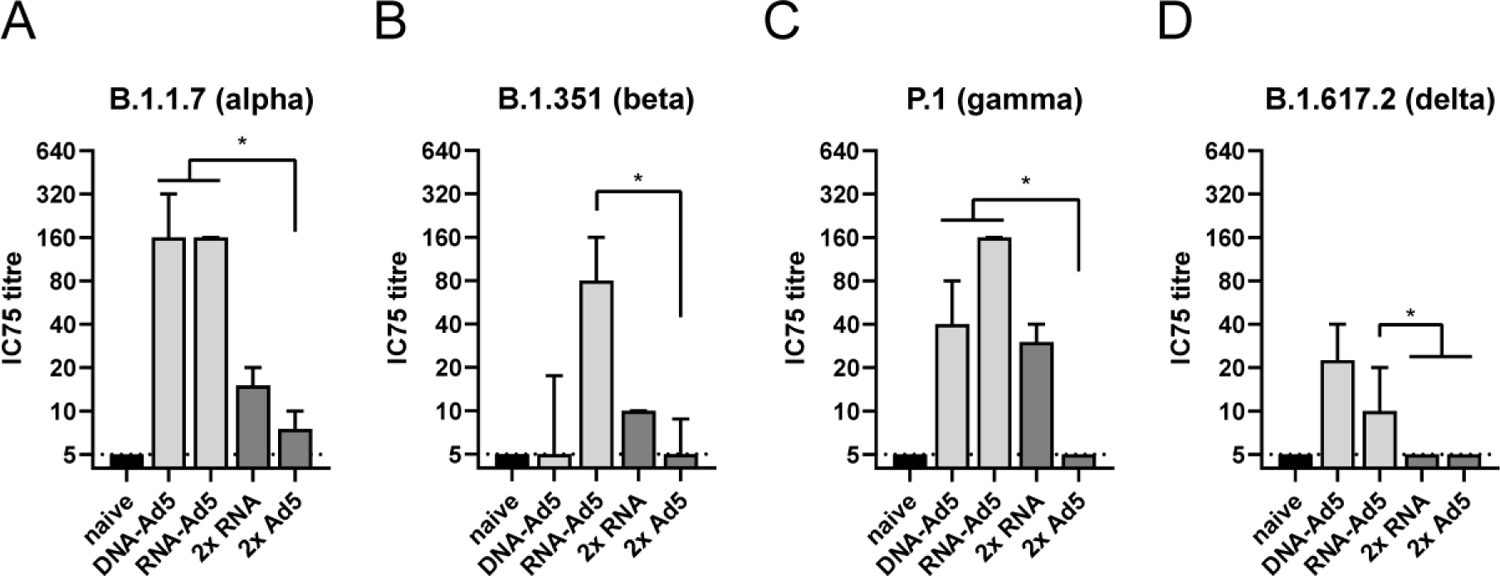
Neutralization of SARS-CoV-2 variants. C57BL/6 mice were vaccinated according to Fig. 5 A. BAL samples were analysed by pseudotype neutralization assays for the neutralization of different variants of concern (A-D). The IC_75_ titre consists of the reciprocal value of the highest dilution able to reduce the infectivity of the respective pseudotype by 75%. Bars represent group medians with interquartile ranges; RNA-Ad5 n=7, other groups n=8. The dashed line indicates the lower limit of detection. Data were analysed by Kruskal-Wallis test (one-way ANOVA) followed by Dunn’s multiple comparison. Statistically significant differences were indicated only among the different vaccine groups (*, p<0.05; **, p<0.005; ***, p<0.0005).

### Lung-resident memory T cells can be efficiently established by a mucosal boost but not by conventional mRNA vaccination

Next, we assessed the induction of systemic and resident T cell memory. Antigen-experienced CD44^+^ CD8^+^ T cells isolated from lung tissue were quantitatively most pronounced in the 2x RNA group (Fig. 7 B). However, by analysing the contribution of tissue-resident (iv-) and vascular (iv+) compartments, a more complex picture emerged. The groups that received two systemic immunizations almost exclusively mounted circulating T cell memory (>95% iv+; Fig. 7 A and B) and consistent to this, the predominant memory phenotypes were T_EFF_, T_EM_, and T_CM_ (Fig. 7 C). CD103^+^CD69^+^ T_RM_ were not established in the lungs of these animals. In complete contrast, the DNA-Ad5 immunized animals displayed mostly T_RM_ but were lacking substantial numbers of circulating memory cells. Importantly, the RNA-Ad5 strategy induced the most comprehensive T cell memory consisting of both circulating subsets and CD103^+^CD69^+^ T cells in the lung.

**Figure 7:**
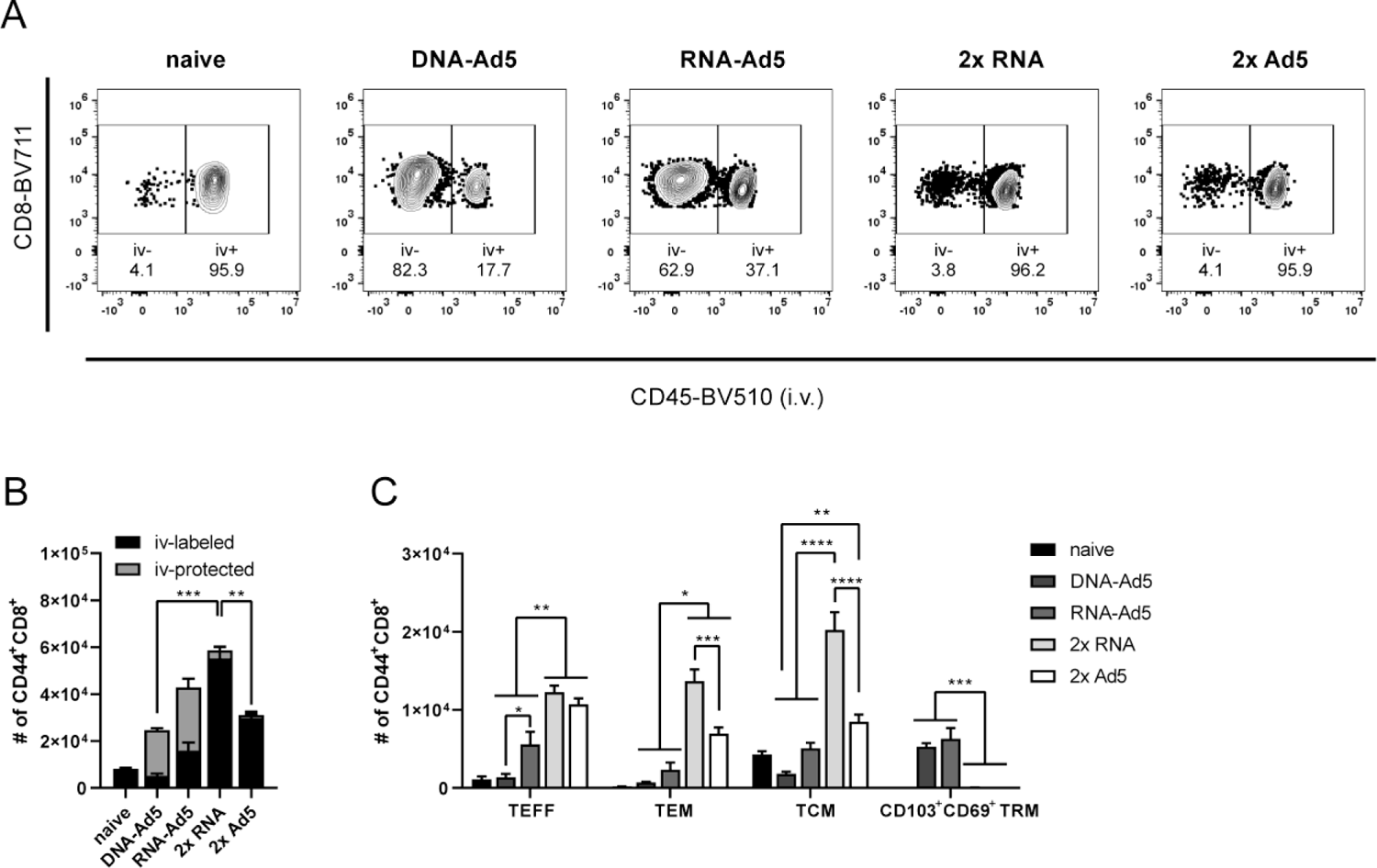
Circulating and tissue-resident memory T cell subsets in the lung. C57BL/6 mice were vaccinated according to Fig. 5 A. Antigen-experienced CD8^+^ T cells were identified by CD44 staining and intravascular staining was used to differentiate between circulating (iv-labelled) and tissue-resident (iv-protected) memory cells. Representative contour plots are shown in (A). (B) The total number of CD44^+^ CD8^+^ with the relative contribution of iv- and iv+ cells are summarized for each group. (C) Within the iv-labelled CD44^+^ CD8^+^ population, effector T cells (T_EFF_; CD127^-^KLRG1^+^), effector memory T cells (T_EM_; CD127^+^KLRG1^+^), and central memory T cells (T_CM_; CD127^+^KLRG1^-^CD69^-^CD103^-^) were defined. Within the iv-protected population, T_RM_ cells were defined as KLRG1^-^CD103^+^CD69^+^. The gating strategy is shown in figure S2. Bars represent group means with SEM; all groups n=4. Data were analysed by one-way ANOVA followed by Tukey’s multiple comparison test. Statistically significant differences were indicated only among the different vaccine groups (*, p<0.05; **, p<0.005; ***, p<0.0005; ****, p<0.0001).

The analysis of spike-specific, cytokine producing CD8^+^ T cells showed a similar compartmentalization. Although the overall numbers of CD107a^+^, IFNγ^+^, and TNFα^+^ CD8^+^ T cells were highest in the lungs of the 2x RNA group, these cells were almost exclusively found in the vascular compartment (iv-labelled, Fig. 8 A-C). The same is true for the homologous immunization with Ad5, albeit reaching much lower percentages of reactive cells. In line with the phenotypic analyses, RNA-Ad5 induced both systemic and local T cell responses, whereas DNA-Ad5 provoked mainly T_RM_. The trends observed for CD8^+^ T cell responses in the iv-labelled lung population were largely mirrored by the splenic responses (Fig. 8 D), further underlining that the former population reflects circulating T cells present in the lung vasculature at the time of sampling. Spike-specific, tissue-resident CD4^+^ T cell responses were also effectively established by the mucosal boost strategies (Fig. 9 A and B) and systemic CD4^+^ T cells in the spleen were induced by all vaccine schedules with two RNA shots being the most effective strategy (Fig. 9 D).

**Figure 8:**
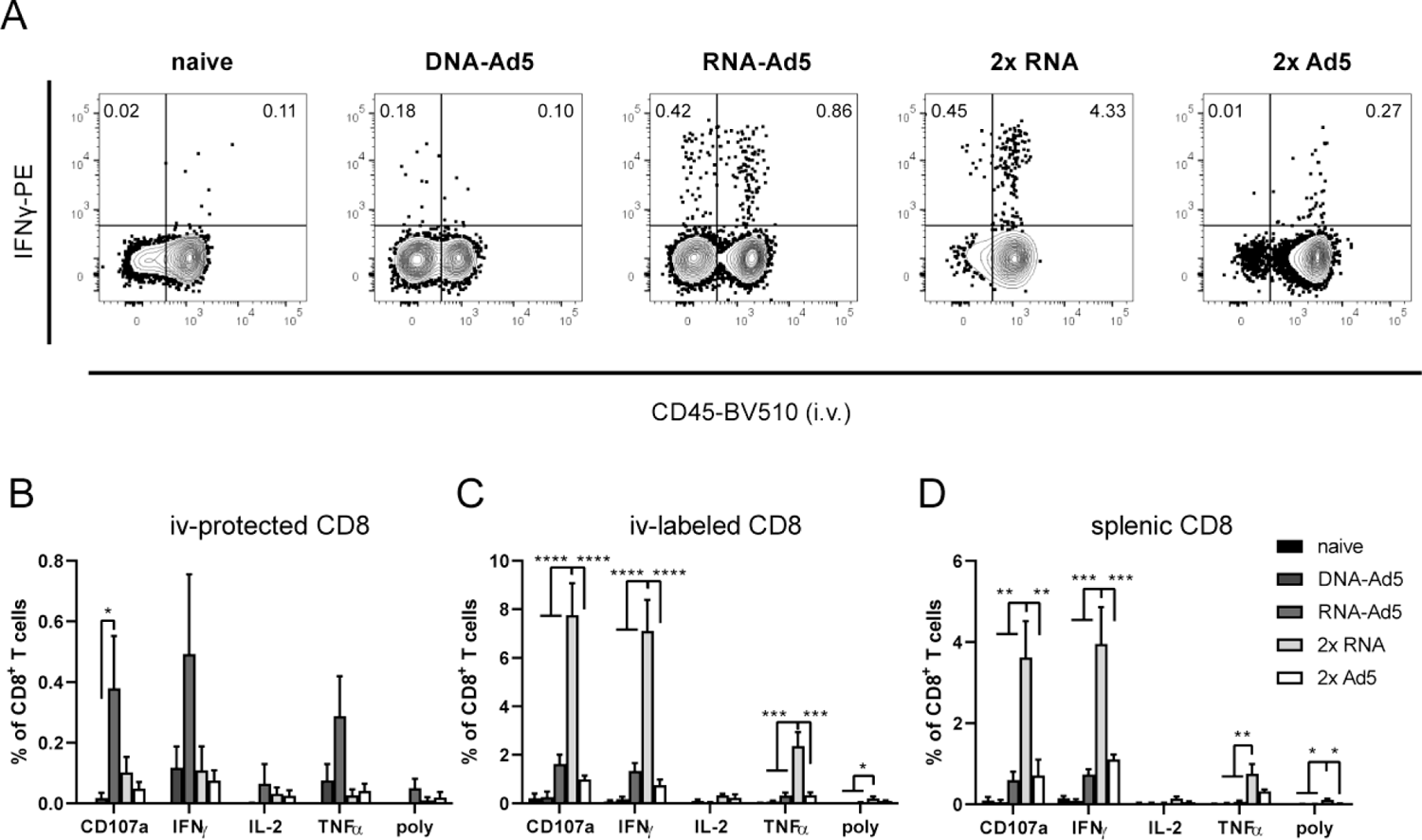
Spike-specific CD8^+^ T cell responses. C57BL/6 mice were vaccinated according to Fig. 5 A. Lung (B and C) and spleen homogenates (D) were restimulated with a peptide pool covering major parts of S and the responding CD8^+^ T cells identified by intracellular staining for accumulated cytokines or staining for CD107a as degranulation marker. (A) Representative contour plots showing IFNγ production in iv+ and iv-lung CD8^+^ T cells. The gating strategy is shown in figure S3. Bars represent group means with SEM; all groups n=4. Data were analysed by one-way ANOVA followed by Tukey’s multiple comparison test. Statistically significant differences were indicated only among the different vaccine groups (*, p<0.05; **, p<0.005; ***, p<0.0005; ****, p<0.0001). poly; polyfunctional T cell population positive for all assessed markers.

**Figure 9:**
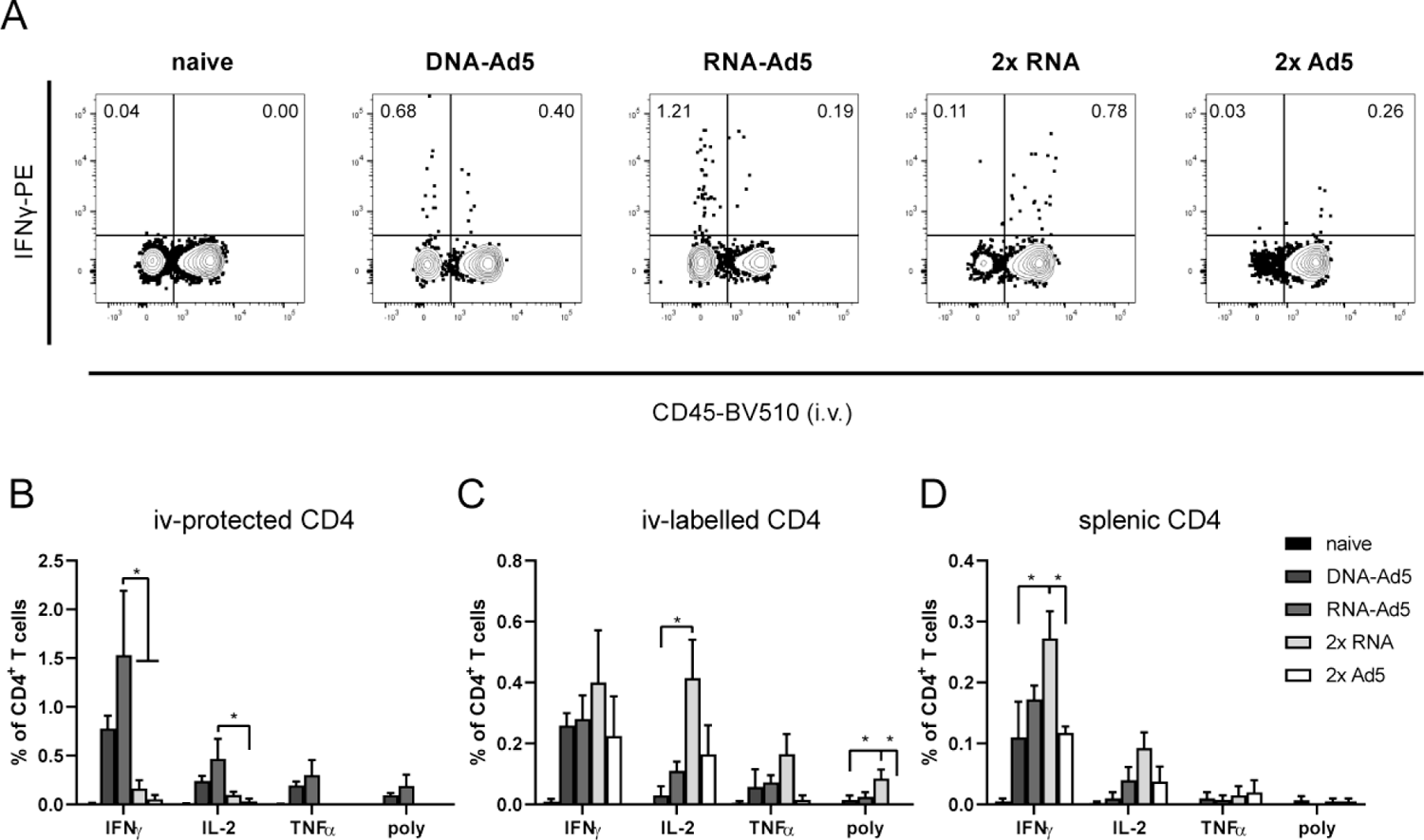
Spike-specific CD4^+^ T cell responses. C57BL/6 mice were vaccinated according to Fig. 5 A. Lung (B and C) and spleen homogenates (D) were restimulated with a peptide pool covering major parts of S and the responding CD4^+^ T cells identified by intracellular staining for accumulated cytokines. (A) Representative contour plots showing IFNγ production in iv+ and iv-lung CD4^+^ T cells. The gating strategy is shown in figure S3. Bars represent group means with SEM; all groups n=4. Data were analysed by one-way ANOVA followed by Tukey’s multiple comparison test. Statistically significant differences were indicated only among the different vaccine groups (*, p<0.05; **, p<0.005; ***, p<0.0005; ****, p<0.0001). poly; polyfunctional T cell population positive for all assessed markers.

In conclusion, only intranasal vaccinations schedules were able to induce profound mucosal immunity in the respiratory tract consisting of neutralizing IgG, IgA, and lung T_RM_. Compared to DNA-Ad5, the RNA-Ad5 strategy provoked a more efficient neutralization of VOCs and established a comprehensive T cell immunity consisting of both T_RM_ and circulatory T cells.

### Systemic and mucosal vaccine schedules effectively protect from experimental SARS-CoV-2 infection

In order to assess the protective efficacy of the vaccination strategies, human ACE2 transgenic mice (K18-hACE2) were immunized as described before and challenged four weeks after the boost immunization with 9×10^3^ FFU of the SARS-CoV-2 strain BavPat1 as previously described ^56^. Since the 2x Ad5 immunization was less immunogenic than the 2x RNA immunization, this group was replaced by another 2x Ad vaccination regime consisting of an intramuscular Ad19a prime followed by the established intranasal Ad5 boost (Fig. 10 A). Seven out of eight unvaccinated control animals reached humane endpoints at day five indicating a severe and lethal course of the disease (Fig. 10 B). They presented weight loss starting at day four post-infection with a concomitant increase of clinical signs (Fig. 10 C and D). In contrast, all vaccinated groups were largely protected from weight loss, clinical signs of disease, and mortality (Fig. 10 B-D). High levels of viral RNA in lung homogenates and BAL fluids were only detected in unvaccinated animals indicating efficient viral replication, while from the vaccinated animals only two of the 2x RNA group had viral RNA copy numbers in the lung above the detection limit (Fig. 10 E). Similarly, infectious virus was retrieved from the lungs of unvaccinated animals but not from the immunized groups (Fig. 10 F). Due to the nature of this challenge model, high viral RNA copy numbers were also detected in the brains of naïve animals (Fig. S7). Although viral RNA was still detectable in the majority of the vaccinated animals, the copy numbers were reduced by 4-5 logs, and no significant differences among the vaccine groups could be seen.

**Figure 10:**
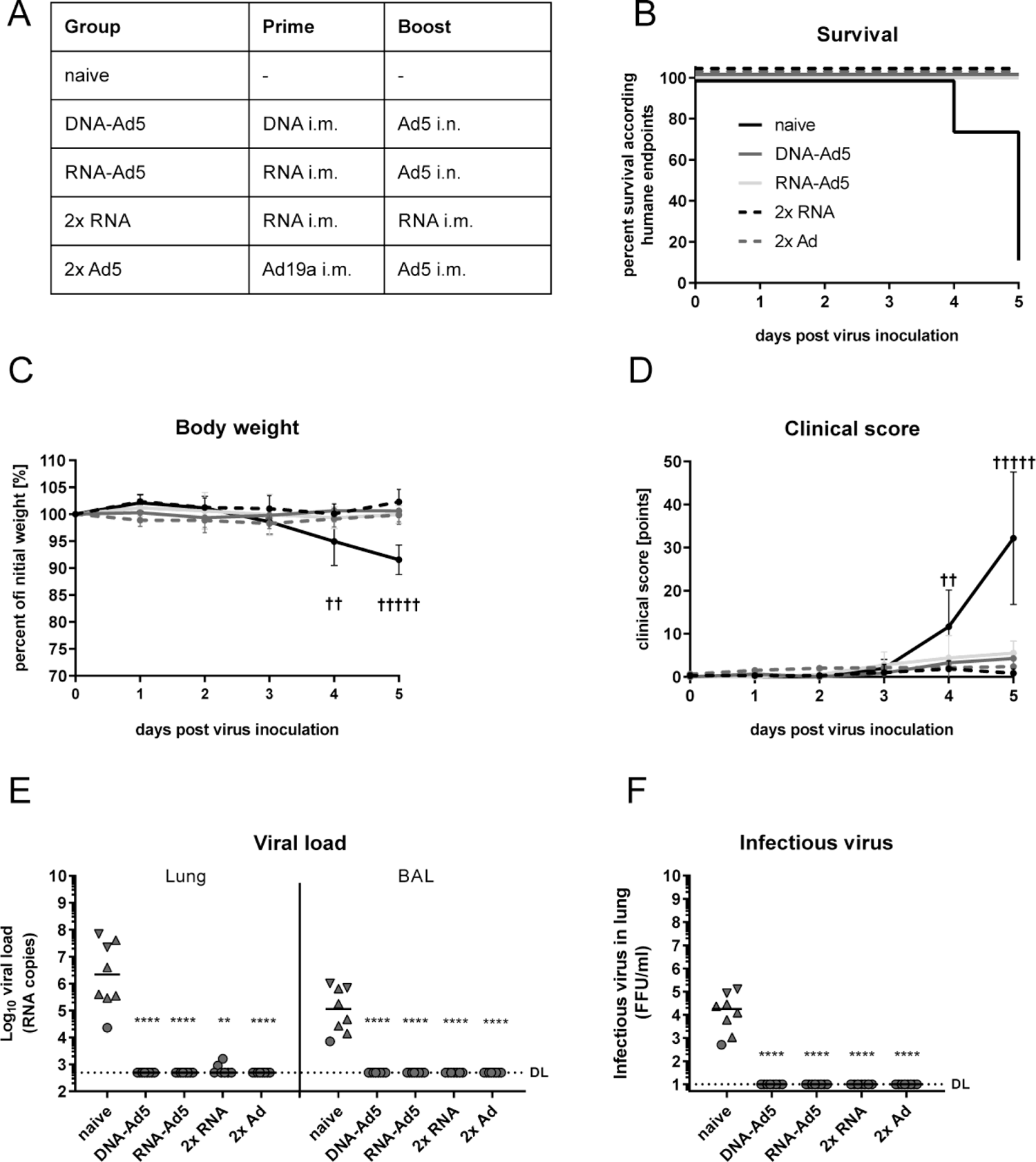
Protective efficacy against SARS-CoV-2 infection. (A) K18-hACE2 mice (2x RNA n=7, other groups n=8) received an intramuscular prime immunization with the spike-encoding DNA (10 µg) followed by electroporation, Ad19-S (10^7^ infectious units), or the mRNA vaccine, Comirnaty® (1 µg). Mice from the heterologous prime-boost groups were boosted four weeks later intranasally or intramuscularly with Ad5-S (10^7^ infectious units). The 2x RNA group received a second dose of mRNA (1 µg). Four weeks after the boost immunization, mice were infected intranasally with 9×10^3^ FFU SARS-CoV-2. All animals were monitored daily for survival (B), body weight (C), and clinical score (D). Animals reaching humane endpoints were euthanized and are marked by a cross (†). Viral RNA copy numbers were assessed in lung homogenates and BAL samples by qRT-PCR (E) and infectious virus was retrieved and titrated from lung homogenates (F). Data points shown represent viral copy number or virus titre of each animal with the median of each group, whereby circles indicate a survival of 5 days post infection and triangles indicates euthanized mouse according humane endpoints at day 4 (▾) or day 5 (▴). The dashed line indicates the lower limit of detection. Data were analysed by Kruskal-Wallis test (one-way ANOVA) and Dunn’s Pairwise Multiple Comparison Procedures as post hoc test in comparison to PBS control (*, p<0.05; **, p<0.005; ***, p<0.0005; ****, p<0.0001).

Taken together, the mucosal boost strategies were able to fully prevent mortality and symptomatic disease upon experimental SARS-CoV-2 infection. The protective efficacy was equal to the current approved vaccination regimen consisting of two intramuscular injections of Comirnaty®.

## Discussion

The SARS-CoV-2 pandemic had and still has a deep impact on social, economic, and healthcare aspects of the world community. As a reaction, academic institutions, biotech companies, and regulatory agencies released safe and effective vaccines in an unprecedented speed. While early in the pandemic the vaccine efficacies of the respective vaccine schedules were in focus, the interest now shifts towards investigating immunogenicity and efficacy of mixed modality vaccinations, the maintenance of long-term immunity, and the protection against emerging variants. So far, the heterologous combination of different vaccine modalities is mostly connected to a superior immunogenicity in preclinical ^57, 58^ and clinical studies ^59–62^. As now most countries with progressed vaccination campaigns discuss the employment of booster vaccinations, a possible next step might be the implementation of mucosal immunizations in order to harness the full potential of mucosal immunity at the entry port of SARS-CoV-2 infections.

To this end, the current study determined the immunogenicity and protective efficacy of mucosal boost vaccination with adenoviral vectors after systemic prime immunizations with a DNA vaccine or an mRNA vaccine that is part of the current vaccine campaigns. The results clearly prove a potent induction of mucosal immune responses by these heterologous strategies not seen after purely systemic immunization schedules. Concomitantly, we observed comparable protective efficacies upon experimental infection among systemic immunization schedules and the heterologous mucosal boost strategies. These results should encourage the exploitation of mucosal booster immunizations as a non-traumatic vaccination modality able to induce strong mucosal immunity in addition to systemic responses. Although not discernible in the present study, this front-line immunity might further inhibit breakthrough infections and transmission risk.

In the first part of the present study, we confirmed that a systemic prime with a DNA vaccine potentiates the immunogenicity of mucosally applied adenoviral vector vaccines ^63^. Most probably, systemic memory cells induced by the priming expand during the recall response and are then recruited to the mucosal site to differentiate into tissue-resident memory cells as reported for T_RM_ in the female reproductive tract ^64^. This is an important finding since it clearly proves the suitability to implement mucosal immunizations into current SARS-CoV-2 vaccination schedules. Although preclinical studies imply significant protection against SARS-CoV-2 in mice, hamsters, ferrets, and non-human primates after one intranasal vaccination with a adenoviral vector vaccines ^47–49, 51, 65^, the first reports from a human clinical trial with an intranasal Ad5-based SARS-CoV-2 vaccine, Altimmune’s AdCOVID, were disappointing and the development was discontinued ^52^. Although safe and well tolerated, the vaccine did not demonstrate sufficient immunogenicity after one or two intranasal doses in previously unvaccinated individuals. The data from a very recent clinical phase I trial underline these findings by providing that serum antibody levels were lower after two intranasal doses of an Ad5-based SARS-CoV-2 vaccine than after one single intramuscular injection ^53^. However, the combination of an mucosal booster immunization with an intramuscular prime resulted in the highest levels of neutralising antibodies reported in that clinical trial. Unfortunately, mucosal immune responses were not assessed. These observations might support our notion of the potential need of pre-existing memory cells to maximize the immunogenicity of an intranasal immunization. While this might seem to complicate the use of nasal vaccines at first sight, one has to keep in mind that mass vaccination campaigns currently employ intramuscular immunizations in large parts of the community. This will finally result in a balanced response between systemic and mucosal immunity.

In most mucosal parameters observed, Ad19a was less immunogenic compared to Ad5, whereas systemic responses, especially CD4^+^ T cell responses, were more efficiently induced. We reported this trend before and speculate that different tropisms of the viral vectors might account for that: Ad5 enters cells via binding to the coxsackie-and-adenovirus receptor (CAR), while Ad19a binds sialic acids and CD46 as entry receptors ^66–69^. Since these molecules are differentially expressed on stromal and immune cells, this might be one aspect explaining the different local and systemic immune profiles.

In the second part of the present study, the impact of the systemic priming modality (RNA/DNA) on the immunogenicity and efficacy of intranasal boost vaccinations with Ad5 was investigated and compared to two systemic immunizations with Ad5 or RNA. Humoral responses in the serum were largely comparable in all groups with the exception that two doses Ad5 provoked weaker responses. It is tempting to speculate that anti-vector immunity induced by the primary immunization might have dampened the effect of the homologous booster. This mechanism is also discussed in the context of the lower vaccine efficacy in humans reported with two standard doses Vaxzevria® (ChAdOx1) compared to the low dose-standard dose schedule ^5^.

Mucosal antibody levels were higher in the groups having received a mucosal boost compared to the repeated systemic vaccination regimens. In regard to the levels of mucosal IgG, this trend was less pronounced as for mucosal IgA levels, presumably because serum IgG can be transudated into the respiratory lumen, whereas IgA is more stringently connected to a local immune reaction. Most importantly, the increased antibody responses in the mucosa also translated into more efficient virus neutralization by BAL samples. Only BAL samples from groups with mucosal vaccinations displayed effective neutralization of all tested VOCs. Although definitive evidence is currently missing, mucosal virus neutralization might be key to supress initial infections with SARS-CoV-2 and therefore minimize the risk of transmission to and by vaccinees. Interestingly, we observed distinct neutralization profiles between the DNA-Ad5 and RNA-Ad5 schemes probably originating from the use of different spike antigens. Thus, it is important to investigate the role of the prefusion conformation stabilization ^70^ regarding neutralization profiles in more detail.

An important advantage of intranasally administered genetic vaccines is the induction of T_RM_ in the respiratory tract. In the present study, tissue-resident memory was exclusively established by mucosal vaccinations. This is congruent with published research showing that local antigen expression is essential for the development of respiratory T_RM_ ^30, 32, 40^. Moreover, in combination with a mucosal boost, a priming immunization with RNA provoked a broader cellular immunity compared to a DNA prime consisting of not only T_RM_ in the lung but also of significant numbers of circulating memory T cells. We speculate that such comprehensive T cell immunity is more effective against breakthrough infections than having only circulating or only resident T cells. Although the chosen challenge model in K18-hACE2 mice did not allow to decipher different degrees of protection, it clearly demonstrates that heterologous prime-boost vaccinations with an intranasal component are at least as protective as the currently licensed vaccine schedules.

To further investigate potential advantages of mucosal immune responses, upcoming studies must mimic the settings more closely that likely contribute to breakthrough infections in vaccinated individuals: age ^71^, comorbidities ^72, 73^, waning immunity, and infection with less neutralization-sensitive variants like B.1.617.2 ^12–18^. However, experimental human challenge studies with a small number of participants might also illuminate this topic similar to challenge studies previously performed in the context of RSV^45, 46^ or Influenza ^74, 75^.

Absolute or mechanistic correlates of protection against SARS-CoV-2 are not yet determined, although neutralizing antibody responses in sera were recently described to be predictive of protection against symptomatic infections ^76, 77^. However, limiting the initial infection rate by mucosal IgA and an early control of viral replication by local CD8^+^ T_RM_ would add another layer of protection, which may be underestimated so far. Furthermore, the rapid inhibition of local replication may result in reduced levels of pro-inflammatory cytokines that partially contribute to tissue damage and severe disease progression ^78^. In face of the encouraging results from the mixed vaccine regimens using RNA vaccine first followed by adenoviral vector vaccines ^59–62^, it might be worthy to utilize an intranasal application for the viral vector boost immunization. This atraumatic, non-invasive application might also reduce the systemic side effects reported for the viral vector vaccines ^79, 80^.

Finally, we demonstrated that the heterologous RNA prime/intranasal Ad5 boost immunization is not inferior to the common gold standard of two intramuscular RNA immunizations in regard to efficacy and additionally results in an unmatched mucosal immune response to SARS-CoV-2. Thus, this study provides the basis to pursue further efficacy studies in non-human primate models or even initiate clinical phase I studies using the currently available vaccines.

## Methods

### Vaccines

Codon-optimized sequences for the N or the spike S protein of SARS-CoV-2 were cloned into the pVAX1 expression plasmid (ThermoFisher) optimized for DNA vaccinations referred to as pVAX1-SARS2-N and pVAX1-SARS2-S ^81^. The encoded S protein is the non-stabilized wildtype protein and based on the sequence of the initial Wuhan isolate (NCBI Reference Sequence: NC_045512.2). Replication-deficient (ΔE1ΔE3) adenoviral vector vaccines based Ad5 or Ad19a/64 encoding the same antigens were produced as previously described ^82^ by Sirion Biotech (Martinsried, Germany). In both vector systems, antigen expression is initiated from a CMV-immediate/early-1-promoter and a bovine BGH polyadenylation signal provides transcription termination. The mRNA vaccine Comirnaty® encodes the stabilized prefusion S protein and is described elsewhere ^83^.

### Immunizations

Six to eight weeks old female BALB/cJRj or C57BL/6 mice were purchased from Janvier (Le Genest-Saint-Isle, France) and housed in individually ventilated cages in accordance with German law and institutional guidelines. The study was approved by an external ethics committee authorized by the Government of Lower Franconia and performed under the project license AZ 55.2.2-2532-2-1179. The research staff was trained in animal care and handling in accordance to the FELASA and GV-SOLAS guidelines. For intramuscular immunizations, inhalative isoflurane anaesthesia was applied and the vaccines were injected in a volume of 30 µl PBS in the *musculus gastrocnemius* of each hind leg. In case of DNA immunizations, the injection was followed by electroporation as described previously ^84^. Under general anaesthesia (100 mg/kg ketamine and 15 mg/kg xylazine), intranasal immunizations were performed by slowly pipetting a volume of 50 µl into one nostril containing the final vaccine dose. Blood was sampled from the retro-orbital sinus under light anaesthesia with inhaled isoflurane. For sampling BAL fluids, mice were killed and the lungs were rinsed twice with 1 ml cold PBS through the cannulated trachea.

### Antigen-specific antibody ELISA

Spike S1, S2, and RBD antibody responses were analysed by ELISA. To this end, ELISA plates were coated with 100 ng of the respective peptide (RBD peptide provided by Diarect GmbH, Freiburg; S1 and S2 peptides kindly provided by Thomas Schumacher from Virion/Serion GmbH, Würzburg) in 100 μl carbonate buffer (50 mM carbonate/bicarbonate, pH 9.6) per well over night at 4°C. Free binding sites were blocked with 5% skimmed milk in PBS-T (PBS containing 0.05% Tween-20) for 1h at RT. BAL samples were diluted in 2% skimmed milk in PBS-T and incubated on the plate for one hour at RT. After three washing steps with 200 μl PBS-T, HRP-coupled anti-mouse IgA (dilution 1:5,000, A90-103P, Bethyl Laboratories) detection antibodies were added for 1h at RT. Subsequently, the plates were washed seven times with PBS-T and after the addition of ECL solution, the signal was measured on a microplate luminometer (VICTOR X5, PerkinElmer).

### FACS-based antibody analysis

A modified version of our previously published serological assay was used ^54^, in which stably transduced HEK 293T cells express the antigen of interest. To analyse quantities of antigen-specific antibodies, 5×10^5^ HEK 293T cells producing SARS-CoV-2 spike or nucleocapsid were incubated for 20 minutes at 4°C with the respective biological sample diluted in 100 µl FACS-PBS (PBS with 0.5% BSA and 1 mM sodium azide) to bind to spike protein on the surface, or in 100 µl permeabilization buffer (0.5% saponin in FACS-PBS) to bind to intracellular nucleocapsid protein. After washing with 200 µl buffer, specific antibodies were bound with polyclonal anti-mouse Ig-FITC (1:300, 4°C, 20 min incubation; BD Biosciences), anti-mouse IgG1-APC (1:300, clone X56), or anti-mouse IgG2a-FITC (1:300, clone R19-15, all BD Biosciences). After further washing, samples were measured on an AttuneNxt (ThermoFisher) and analysed using FlowJo software (Tree Star Inc.).

### Virus neutralization assay

Serial dilutions of sera and BALs were incubated with 2000 PFU of an early SARS-CoV-2 isolate (GISAID EPI ISL 406862 Germany/BavPat1/2020) in 100 µl OptiPro medium supplemented with 1x GlutaMAX (both Gibco) for 1h at 37°C. Subsequently, the mixture was added onto a confluent monolayer of Vero E6 cells (seeded the day before at 10^4^ cells per well in a 96-well plate). After 1h, the mixture was removed from the cells and 100 µl OptiPro medium supplemented with 1x GlutaMAX (both Gibco) was added. After 24h incubation at 37°C and 5% CO_2_, cells were fixed with 100 µl 4% paraformaldehyde for 20 min at RT and permeabilized with 100 µl 0.5% Triton X-100 in PBS for 15min at RT. After a blocking step with 100 µl 5% skimmed milk in PBS for 1h at RT, cells were stained with purified immunoglobulins from a SARS-CoV-2 convalescent patient in 2% skimmed milk for 1h at 4°C. After three washing steps with 200 µl PBS, 100 µl of anti-human IgG FITC (1:200, 109-096-003, Jackson Immunoresearch) were added diluted in 2% skimmed milk and incubated for 1h at 4°C in the dark. After another three washing steps with 200 µl PBS, plaques were counted with an ELISPOT reader (Cellular Technology limited BioSpot). Infected wells without serum were used as reference to determine the 75% plaque reduction neutralization titre (PRNT75).

### Pseudotype neutralization assay

Neutralization of various spike variants was assessed with the help of spike-pseudotyped simian immunodeficiency virus particles as described before ^85^. For the production of pseudotyped reporter particles, HEK293T cells were transfected with the SIV-based self-inactivating vector encoding luciferase (pGAE-LucW), the SIV-based packaging plasmid (pAdSIV3), and the respective spike variant-encoding plasmid ^86–88^. For this purpose, 2×10^7^ HEK293T cells were seeded the day before in a 175 cm^2^ flask in Dulbecco’s Modified Eagle’s Medium (DMEM; Gibco) containing 10% FCS, 2 mM L-Glutamine, and 100 units/ml penicillin/streptomycin (D10 medium). The transfection mixture was prepared by mixing 20 µg of each plasmid with 180 µg polyethylenimine in 5 ml DMEM without additives. 15 min later, the mixture was added to the cells. After 4-8h incubation, the medium was exchanged to 25 ml DMEM containing 1.5% FCS, 2 mM L-Glutamine, and 100 units/ml penicillin/streptomycin. 72 h post-transfection, the supernatants containing the lentiviral particles were harvested, sterile filtrated (0.45 µm membrane), and stored at −20°C. HEK293T-ACE2 cells stably expressing the human Angiotensin-converting enzyme 2 (ACE2) were transduced with the dilutions of the pseudotypes. The amount of lentiviral particles resulting in luciferase signals of 2-10 ×10^4^ RLU/s were used for the latter assay.

For the assessment of pseudotype neutralization, HEK293T-ACE2 cells were seeded at 2×10^4^ cells/well in 100 µl D10 in a 96well flat bottom plate. 24h later, 60 µl of serial dilutions of the BAL samples were incubated with 60 µl lentiviral particles for 1 h at 37°C. HEK293T cells were washed once with PBS and the particle-sample mix was added to the cells. 48 h later, medium was discarded and the cells lysed with 100 µl Bright Glo lysis buffer (Promega) for 15 min at 37°C. Three minutes later, after the addition of 25 µl Bright Glo substrate (Promega), the luciferase signal was assessed on a microplate luminometer (VICTOR X5, PerkinElmer). Neutralization titres are determined as the last reciprocal dilution that inhibits more than 75% of the luciferase signal measured in positive controls (inhibitors concentration 75%, IC75).

### T cell assays

For the definition of circulatory and tissue-resident T cells, mice were injected with 3 µg anti-CD45-BV510 (clone 30-F11, Biolegend) intravenously and were euthanized three minutes later with an overdose of inhaled isoflurane. Spleens and lungs were harvested. The latter ones were cut into small pieces followed by incubation for 45 min at 37 °C with 500 units Collagenase D and 160 units DNase I in 2 ml R10 medium (RPMI 1640 supplemented with 10 % FCS, 2 mM L-Glutamine, 10 mM HEPES, 50 μM β-mercaptoethanol and 1 % penicillin/streptomycin). Digested lung tissues and spleens were mashed through a 70 µm cell strainer before the single cell suspensions were subjected to an ammonium-chloride-potassium lysis. One million splenocytes or 20% of the total lung cell suspension were plated per well in a 96-well round-bottom plate for in vitro restimulation and phenotype assays.

For the restimulation, samples were incubated for 6 hours (or 24 hours in case of Fig. 8 und 9) in 200 µl R10 medium containing monensin (2 µM), anti-CD28 (1 µg/ml, eBioscience), anti-CD107a-FITC (clone eBio1D4B, eBioscience), and the respective SARS-CoV-2 peptide pool (0.6 nmol/peptide, S and N pools from Miltenyi Biotec, 130-126-701 and 130-126-699). Unstimulated samples were used for subtraction of background cytokine production. Cells were stained after the stimulation with anti-CD8a-Pacific blue (1:300, clone 53-6.7, BD Biosciences), anti-CD4-PerCP (1:2000, clone RM4-5, eBioscience) and Fixable Viability Dye eFluor® 780 (1:4000, eBioscience) in FACS-PBS for 20 min at 4°C. After fixation (2% paraformaldehyde, 20 min, 4°C) and permeabilization (0.5% saponin in FACS-PBS, 10 min, 4°C), cells were stained intracellularly with anti-IL-2-APC (1:300, clone JES6-5H4, BD Biosciences), anti-TNFα-PECy7 (1:300, clone MPG-XT22, BD Biosciences), and anti-IFNy-PE (1:300, clone XMG1.2, eBioscience). The gating strategy is shown in Fig. S3.

For the phenotype analyses, cells were stained in FACS-PBS with anti-CD8-BV711 (1:300, clone 53-6.7, BioLegend), anti-CD4-SB600 (1:1000, clone RM4-5, BioLegend), anti-CD127-FITC (1:500, clone A7R34, BioLegend), anti-CD69-PerCP-Cy5.5 (1:300, clone H1.2F3, BioLegend), anti-CD103-PE (1:200, clone 2E7, Invitrogen), anti-KLRG1-PE-Cy7 (1:300, clone 2F1, Invitrogen), anti-CD44-APC (1:5000, clone IM7, BioLegend), and Fixable Viability Dye eFluor® 780 (1:4000, eBioscience). Data were acquired on an AttuneNxt (ThermoFisher) or on a LSRII (BD Biosciences) and analysed using FlowJo^TM^ software (Tree Star Inc.). The gating strategy is shown in Fig. S2.

### SARS-CoV-2 infection model

The infection experiments were approved by local authorities after review by an ethical commission (TVV 21/20). Eleven weeks old, female K18-hACE2 mice (Jackson Laboratory, Bar Harbor, USA) were immunized as described before and infected four weeks after the boost immunization intranasally with 9×10^3^ focus-forming units (FFU) of the SARS-CoV-2 strain BavPat1 in a total volume of 50 µl under light anaesthesia with inhaled isoflurane. Animals were monitored daily for body weight and clinical score. The following parameters were evaluated in the scoring system: weight loss and body posture (0-20 points), general conditions including the appearance of fur and eye closure (0-20 points), reduced activity and general behaviour changes (0-20 points), and limb paralysis (0-20 points). Mice were euthanized at day 5 after infection or earlier if a cumulative clinical score of 20 or more was reached. After euthanasia, the lungs were filled with 800 µl PBS and the left lung was tied off. The BAL of the right lung was taken and repeated with two more washes each with 400 µl. The right lungs as well as the right hemispheres of the brains were homogenized in 1 ml PBS using a gentleMACS Octo Dissociator (Miltenyi Biotec) and viral RNA was isolated from 140 µl cleared homogenate or BAL fluid using QIAamp Viral RNA Mini Kit (Qiagen). RT-qPCR reactions were performed using TaqMan® Fast Virus 1-Step Master Mix (ThermoFisher) and 5 µl of isolated RNA as a template to detect a 132 bp sequence in the ORF1b/NSP14. Primer and probe sequences were as follows: forward primer, 3’-TGGGGYTTTACRGGTAACCT-5’; reverse primer, AACRCGCTTAACAAAGCACTC; probe, 3’-FAM-TAGTTGTGATGCWATCATGACTAG-TAMRA-5’. Synthetic SARS-CoV-2-RNA (Twist Bioscience) was used as a quantitative standard to obtain viral copy numbers ^89^. For the detection of infectious virus in BAL and the lung, Vero E6 cells were seeded at 2×10^4^ cells/well in a 96-well plate in 200 µl of D10 for confluent monolayer 24 h prior to infection. After medium change to D10, a two-fold-serial dilution of BALs or lung homogenates were applied to the cells for 3 hours. After replacing the supernatant with overlay medium (DMEM with 1 % methyl cellulose, 2 % FBS and 1% penicillin/streptomycin), cells were incubated for further 27 hours. SARS-CoV-2 infected cells were visualized using SARS-CoV-2 S-protein specific immunochemistry staining with anti-SARS-CoV-2 spike glycoprotein S1 antibody (Abcam) ^90^.

## Statistical analyses

Results are shown as mean ± SEM or as median ± interquartile range except it is described differently. Statistical analyses were performed with Prism 8.0 (GraphPad Software, Inc.). A p value of <0.05 was considered to be statistically significant. For reasons of clarity, significances are only shown among the vaccine groups.

## Data availability

All data are included in the manuscript or in the supplementary material.

## Acknowledgements

We would like to thank Drew Hannaman (Ichor Medical Systems, Inc.) for providing the TriGrid electrode array for DNA electroporation, Anne-Kathrin Donner for excellent technical assistance, and the group of Dr. Jasmin Fertey (Fraunhofer IZI, Leipzig, Germany) for providing viral stocks for mouse infections. Moreover, we kindly thank Thomas Schumacher (Virion/Serion GmbH, Würzburg, Germany) and Diarect GmbH (Freiburg, Germany) for providing the Spike peptides. The ELISPOT Analyzer was obtained with financial support from Fondation Dormeur, Vaduz. This work was supported within the FOR-COVID project funded by the Bavarian State ministry for Science and the Arts. Further support was provided by B-FAST, a BMBF-funded project of the Netzwerk Universitätsmedizin (NaFoUniMedCovid19; FKZ: 01KX2021) and funds from the Deutsche Forschungsgemeinschaft (DFG) through the research training group RTG 2504 (project number: 401821119). S.P. acknowledges funding by BMBF (01KI2006D, 01KI20328A, 01KI20396, 01KX2021), the Ministry for Science and Culture of Lower Saxony (14-76103-184, MWK HZI COVID-19) DFG (PO 716/11-1, PO 716/14-1).

## Author contributions

D.L., T.G., K.Ü., and M.T. conceived and designed the study. D.L., J.F., J.W., A.V.A., V.E., N.U., L.I., A.S., F.O., A.S.P., P.I., and K.F. collected the data. D.L., J.F., J.W., A.V.A., V.E., N.U., L.I., T.G., and M.T. performed the analysis. S.M.-S., A.C., A.E., M.H., S.P., C.P., T.W., and C.T. contributed critical reagents. D.L. and M.T. drafted the manuscript, which was then critically reviewed and approved by all co-authors.

