## Supplemental Appendix for "Protective mucosal immunity against SARS-CoV-2 after heterologous systemic RNA-mucosal adenoviral vector immunization"

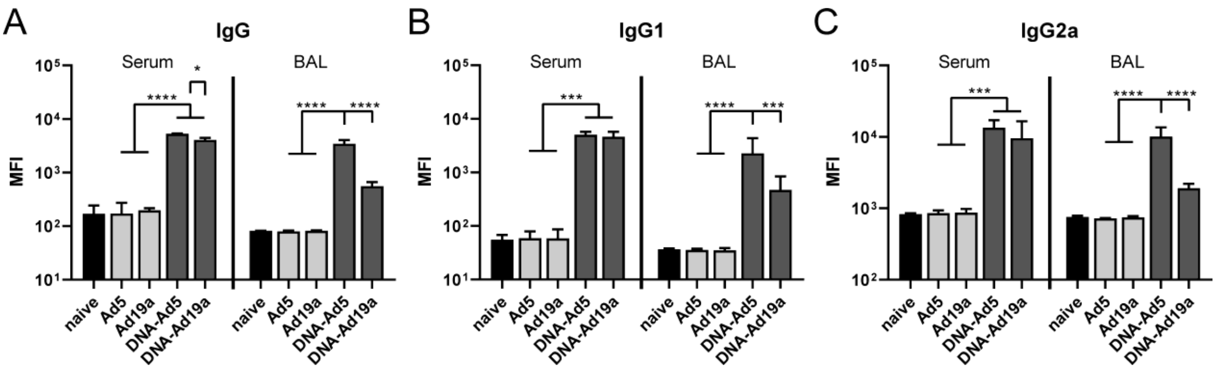


**Figure S1: Nucleocapsid-directed humoral response after intranasal immunization with Ad5- or Ad19a-based viral vector vaccines.** BALB/c mice were vaccinated according to Fig. 1 A. Nucleocapsid-specific IgG (A), IgG1 (B), and IgG2a (C) were assessed by a flow cytometric approach (dilutions: Sera 1:400, BAL 1:100. Bars represent group medians with interquartile ranges; naïve n=4; DNA-Ad5 n=5; other groups n=6. Data were analysed by one-way ANOVA followed by Tukey's post test. Statistically significant differences were indicated only among the different vaccine groups (*, p<0.05; **, p<0.005; ***, p<0.0005; ****, p<0.0001).

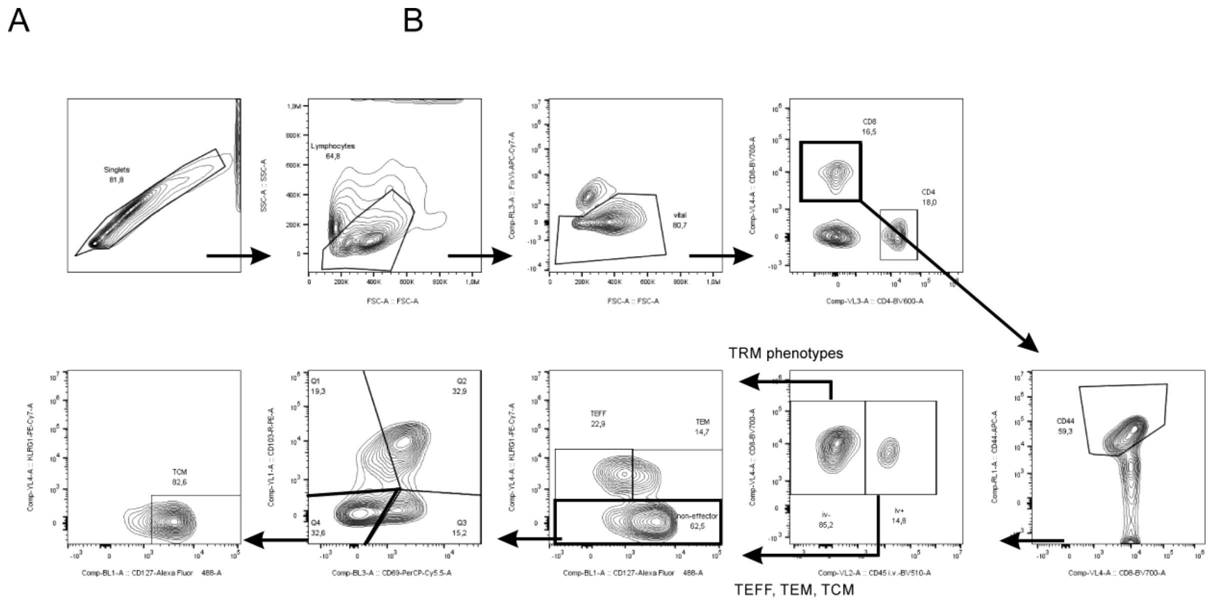


**Figure S2: Gating strategy for circulatory and tissue-resident memory populations.** Depicted is the gating path in a lung sample from a DNA-Ad5 immunized animal. Memory phenotypes are pre-gated on iv+ or iv- cells for circulatory or tissue-resident populations, respectively.

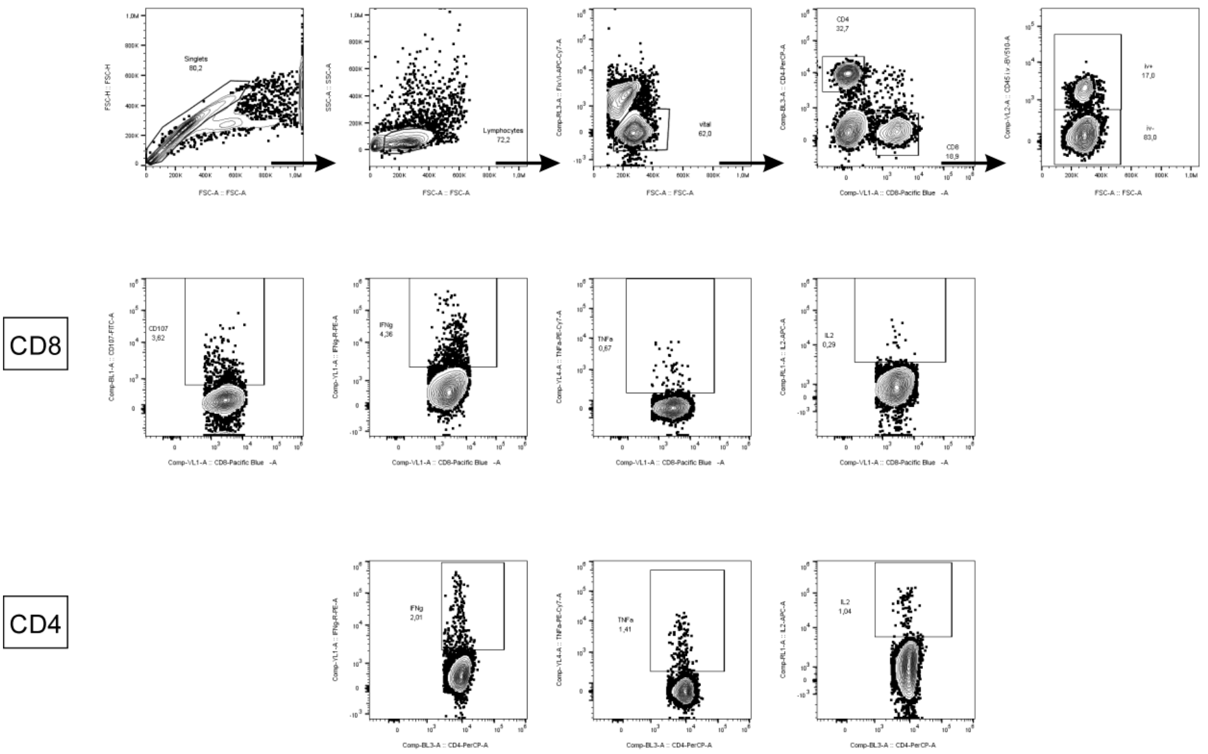


**Figure S3: Gating strategy for antigen-specific T cell responses.** Depicted is the gating path in a lung sample from a DNA-Ad5 immunized animal. Depending on the respective experimental part, antigen-specific T cells were analysed in the total, iv-, or iv+ T cell populations. CD107a as a marker for recent degranulation was only assessed in $CD8^+$ T cells.

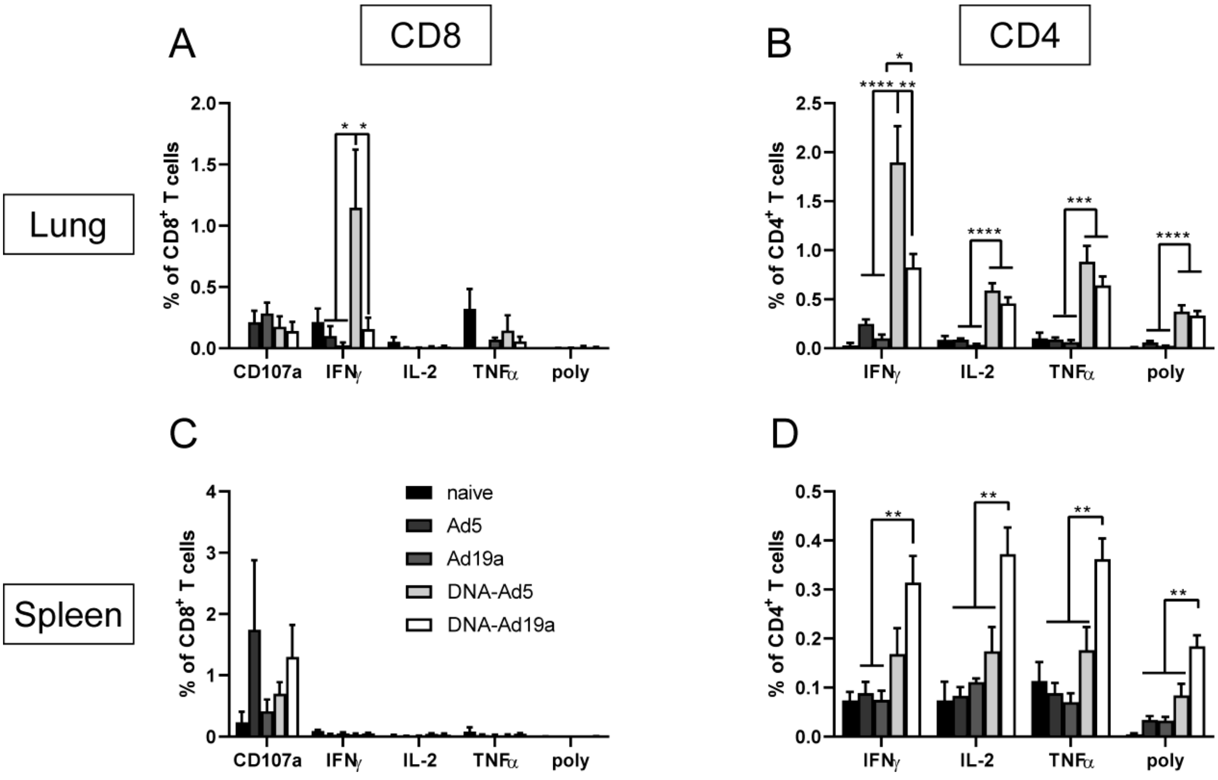


**Figure S4: Nucleocapsid-specific T cell responses after intranasal immunization with Ad5- or Ad19a-based viral vector vaccines.** BALB/c mice were vaccinated according to Fig. 1 A. Lung and spleen homogenates were restimulated with peptide pools covering the complete SARS-CoV-2 N and the responding CD8$^+$ (A and C) and CD4$^+$ T cells (B and D) were identified by intracellular staining for accumulated cytokines or staining for CD107a as degranulation marker. Bars represent group means with SEM; naïve n=4; DNA-Ad5 n=5; other groups n=6. The gating strategy is shown in figure S3. Data were analysed by one-way ANOVA followed by Tukey's multiple comparison test. Statistically significant differences were indicated only among the different vaccine groups (*, p<0.05; **, p<0.005; ***, p<0.0005; ****, p<0.0001). poly; polyfunctional T cell population positive for all assessed markers.

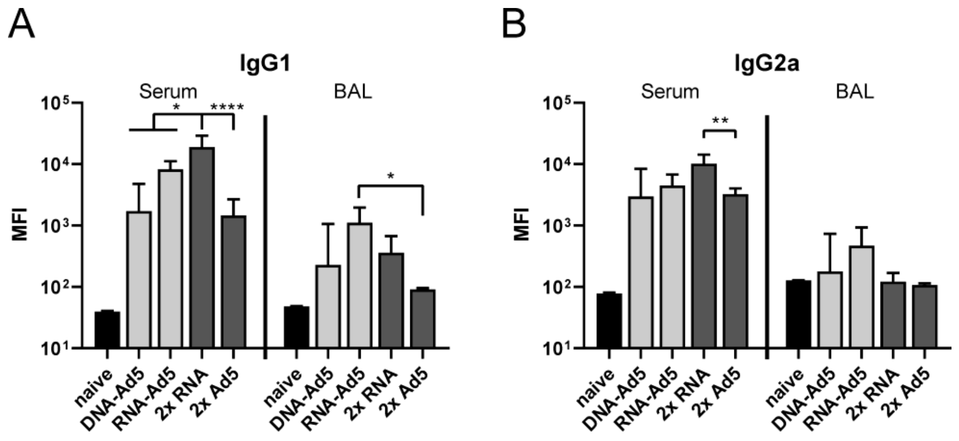


**Figure S5: IgG1 and IgG2 responses after boost immunization.** C57BL/6 mice were vaccinated according to Fig. 5 A. Spike-specific IgG1 (A) and IgG2a (B) in sera and BALs were assessed by a flow cytometric approach (Sera 1:800, BAL 1:20). Bars represent group medians with interquartile ranges; sera all groups n=8; BALs RNA-Ad5 n=7, other groups n=8. Data were analysed by one-way ANOVA followed by Tukey's post test. Statistical significant differences were indicated only among the different vaccine groups (*, p<0.05; **, p<0.005; ***, p<0.0005; ****, p<0.0001).

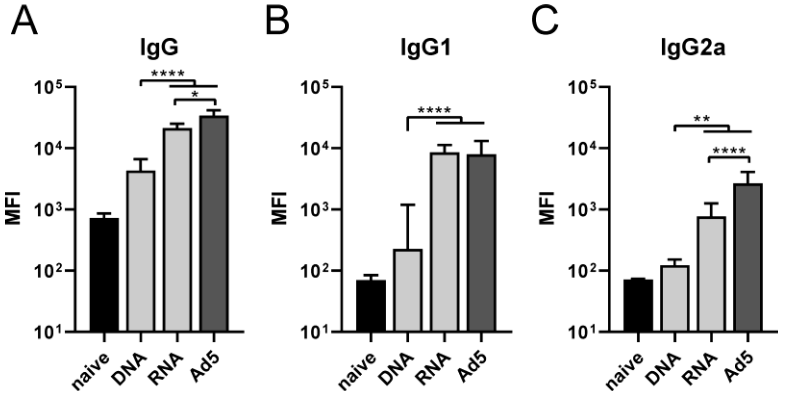


**Figure S6: Humoral responses after prime immunization.** Blood samples were taken 27 days after the prime immunizations and spike-specific IgG (A), IgG1 (B), and IgG2a (C) were assessed by a flow cytometric approach (dilution: 1:200). Bars represent group medians with interquartile ranges; naive n=8, DNA n=16, RNA n=24, Ad5 n=7. Data were analysed by one-way ANOVA followed by Tukey's post test. Statistically significant differences were indicated only among the different vaccine groups (*, p<0.05; **, p<0.005; ***, p<0.0005; ****, p<0.0001).

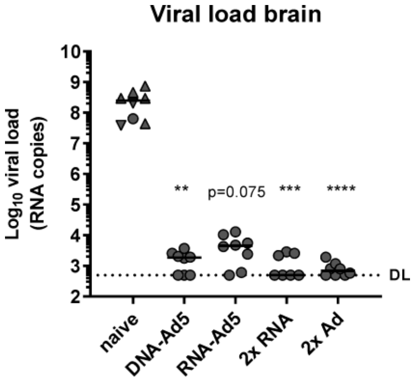


**Figure S7: Virus RNA in brain tissue.** Viral RNA copy numbers were assessed in brain homogenates by qRT-PCR. Data points shown represent viral copy number of each animal with the median of each group, whereby circles indicate a survival of 5 days post infection and triangles indicates euthanized mouse according humane endpoints at day 4 (▼) or day 5 (▲). The dashed line indicates the lower limit of detection. Data were analysed by Kruskal-Wallis test (one-way ANOVA) and Dunn's Pairwise Multiple Comparison Procedures as post hoc test in comparison to PBS control (*, $p<0.05$; **, $p<0.005$; ***, $p<0.0005$; ****, $p<0.0001$).
